# Selection against maternal microRNA target sites in maternal transcripts

**DOI:** 10.1101/012757

**Authors:** Antonio Marco

**Author notes:** Corresponding author: Antonio Marco, School of Biological Sciences, University of Essex, Wivenhoe Park, Colchester CO4 3SQ, United Kingdom.

## Abstract

In animals, before the zygotic genome is expressed, the egg already contains gene products deposited by the mother. These maternal products are crucial during the initial steps of development. In *Drosophila melanogaster* a large number of maternal products are found in the oocyte, some of which are indispensable. Many of these products are RNA molecules, such as gene transcripts and ribosomal RNAs. Recently, microRNAs – small RNA gene regulators – have been detected early during development and are important in these initial steps. The presence of some microRNAs in unfertilized eggs has been reported, but whether they have a functional impact in the egg or early embryo has not being explored. I have extracted and sequenced small RNAs from *Drosophila* unfertilized eggs. The unfertilized egg is rich in small RNAs and contains multiple microRNA products. Maternal microRNAs are often encoded within the intron of maternal genes, suggesting that many maternal microRNAs are the product of transcriptional hitch-hiking. Comparative genomics analyses suggest that maternal transcripts tend to avoid target sites for maternal microRNAs. I also developed a microRNA target mutation model to study the functional impact of polymorphisms at microRNA target sites. The analysis of *Drosophila* populations suggests that there is selection against maternal microRNA target sites in maternal transcripts. A potential role of the maternal microRNA mir-9c in maternal-to-zygotic transition is also discussed. In conclusion, maternal microRNAs in *Drosophila* have a functional impact in maternal protein-coding transcripts.

## INTRODUCTION

In animals, the initial steps of embryonic development are driven by the gene products deposited by the mother into the egg. For instance, in *Drosophila melanogaster*, the anteroposterior axis is determined by the presence of maternal transcripts from genes such as *bicoid* and *nanos* (Lawrence 1992). Recently, the role of microRNAs during development have become a major research area. MicroRNAs are small RNA molecules that regulate gene expression by targeting gene transcripts by sequence complementarity. MicroRNAs are expressed during early development (Aravin *et al.* 2003; Aboobaker *et al.* 2005), and they target other embryonic expressed gene transcripts (Enright *et al.* 2003; Lai *et al.* 2003). As a matter of fact, a number of homeotic genes detected by genetic analysis were later shown to be microRNA encoding genes (reviewed in (Marco 2012)). Traditionally, maternal genes have been identified by genetic analysis (Lawrence 1992). However, the characterization of maternal microRNAs is particularly difficult as they are too short for standard genetic analyses. Thanks to the development of high-throughput technologies such as RNAseq and microarrays, it is now possible to isolate small RNAs directly from egg extracts. For instance, the microRNA content of mouse (Tang *et al.* 2007) and cow (Tesfaye *et al.* 2009) oocytes have been characterized with this high-throughput approach. In other cases, such as in zebrafish (Chen *et al.* 2005) and *Xenopus* (Watanabe *et al.* 2005), microRNAs appear to have a minor presence in oocytes.

Several lines of evidence suggested that, in *Drosophila*, maternally transmitted microRNAs are important. First, some microRNAs are highly abundant during early development (Ruby *et al.* 2007). Also, the enzymes responsible for microRNA biogenesis are present in the ovaries (Robinson *et al.* 2013) and microRNAs may have a role in oocyte maturation (Nakahara *et al.* 2005). Indeed, mature microRNAs have been identified in *Drosophila* unfertilized eggs (Lee *et al.* 2004, 2014; Votruba 2009). Recently, it has been shown that maternally transmitted microRNAs are adenylated during the maternal-to-zygotic transition (MZT) (Lee *et al.* 2014). Whether maternal microRNAs have a functional impact in *Drosophila* eggs is still unknown. To identify which microRNAs are maternally transmitted I extracted and sequenced small RNAs from *Drosophila* unfertilized eggs. To explore their potential function I predicted their targets in maternal and zygotic gene products. The evolutionary impact of maternal microRNAs was estimated by using comparative genomics and population genetics.

## MATERIALS AND METHODS

**Flies and egg collection:** Fly stocks used in this study, with Bloomington reference number in square brackets, were: w^1118^ [#3605] and Oregon-R-modENCODE [#25221]. All flies were kept at 25°C on cornmeal based media, with 12 hours light/dark cycles. Virgin females were sorted at the pupae stage to avoid any unwanted fertilization. (Previous attempts selecting for <6 hours females produced a small yet significant number of fertilized eggs.) In a population cage I let 80-100 females to lay eggs in apple juice agar plates for 8 hours, collecting 1 hour after dawn. Eggs were collected with a sieve and washed with saline solution. Eggs from virgin females do not degenerate even several hours after laying (Tsien and Wattiaux 1971).

**RNA extraction, sequencing and profiling:** Total RNA was extracted from eggs or early embryos with TRIzol reagent (Life Technologies), following instructions given by the manufacturer, and dissolving the RNA in RNase-free water. For RNA sequencing, a cDNA library was generated with TruSeq Small RNA Sample Preparation Kit (Illumina). Amplified cDNA constructs were size selected in a 6% PAGE gel for 145 to 160 bp (fragments including RNA-derived sequences of size ∼20-30 bp plus adapters). Size selected cDNAs were purified and precipitated with ethanol, and DNA integrity was checked with TapeStation (Agilent). Samples were sequenced with Illumina MiSeq in the Genomics Core Facility at the University of Manchester. A total of 4,507,291 reads were sequenced, most of them (95.5%) deriving from ribosomal RNAs which is expected in *Drosophila* where the majority of small RNAs are 2S rRNA (Seitz *et al.* 2008). 13,114 reads were identified as microRNA products. Sequence reads are available from Gene Expression Omnibus (GEO) at NCBI (accession number: GSE63488).

Illumina MiSeq produces 50 bp sequence reads. Hence, I removed adapters with Cutadapt (https://code.google.com/p/cutadapt/) and mapped the processed reads of size 18-26 bp to known microRNAs from miRBase v.20 (Kozomara and Griffiths-Jones 2014), using Bowtie v.0.12.7 (Langmead *et al.* 2009), allowing no mismatches and considering reads mapping to up to five positions. Other RNA collections from embryos and ovaries were also analysed: 0-1h embryos, 2-6h embryos, 6-10 h embryos (Ruby *et al.* 2006) and ovaries (Czech *et al.* 2008). Expression profiling in Figure 1 was done with R (R Development Core Team 2004), scaling the Z-scores of the heatmap across rows, and generating a hierarchical tree of microRNAs with complete linkage clustering.

**Figure 1.**
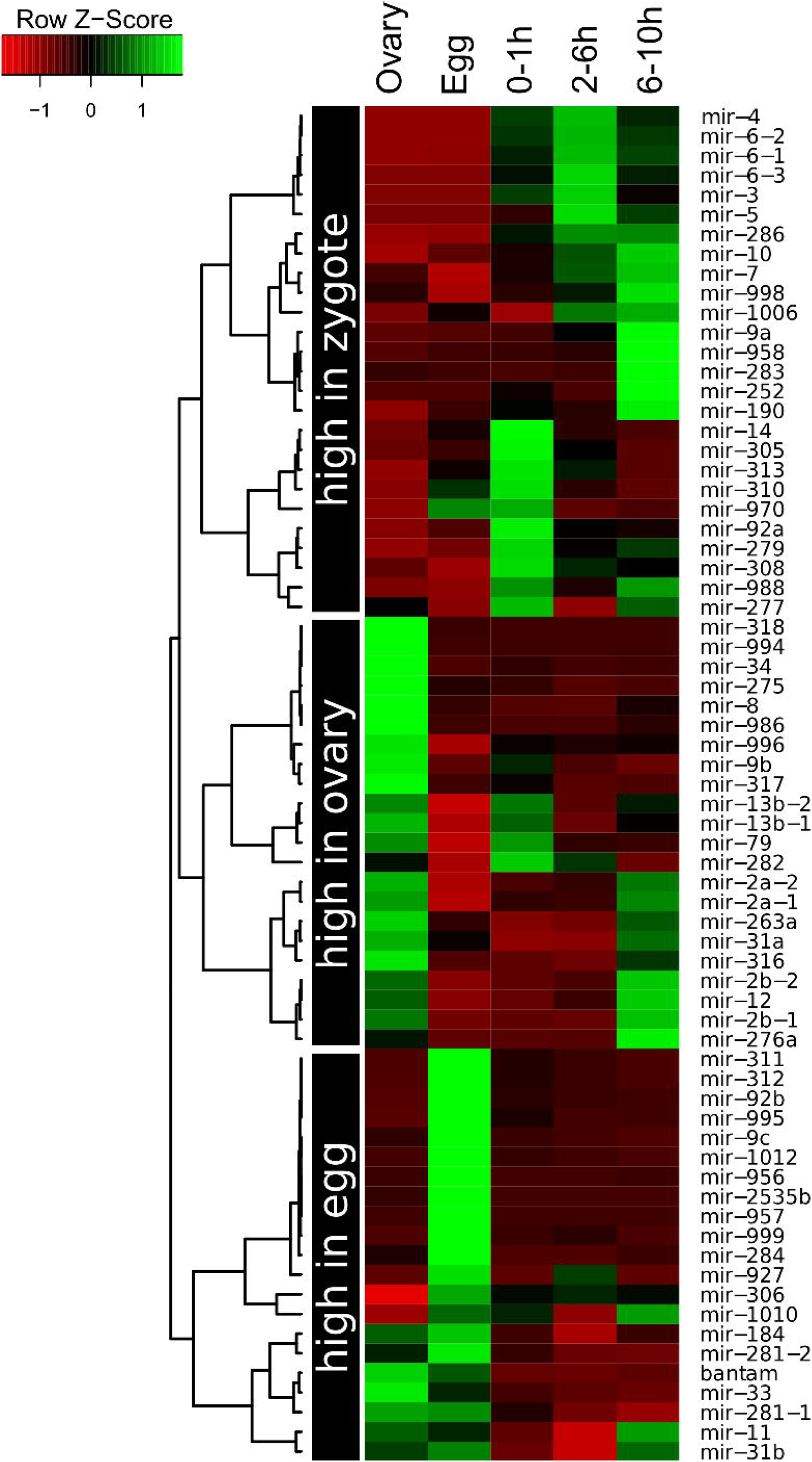
Expression profile of maternal microRNAs in *Drosophila melanogaster*. The hierarchical tree is split into three categories of microRNAs: high abundance in ovaries compared to the other stages; those which are mainly present in the unfertilized eggs and those which have a higher expression level later during development.

The presence in eggs of mature microRNAs was validated with Mir-X first-strand synthesis and SYBR qRT-PCR assays manufactured by Clontech Laboratories, Inc. MicroRNA cDNA libraries were constructed for unfertilized eggs and 2-6 hours old embryos from Oregon-R flies, following the indications from the manufacturer. Primers for microRNA-specific amplification during qPCRs were: let-7-5p (5'-TGAGGTAGTAGGTTGTATAGT-3'), miR-34-5p (5'-TGGCAGTGTGGTTAGCTGGTTGTG-3'), miR-311-3p (5'-TATTGCACATTCACCGGCCTGA-3'), mir-92b-3p (5'-AATTGCACTAGTCCCGGCCTGC-3'), miR-184-3p (5'-TGGACGGAGAACTGATAAGGGC-3'), miR-9c-5p (5'-TCTTTGGTATTCTAGCTGTAGA-3'), bantam-3p (5'-TGAGATCATTTTGAAAGCTGATT-3'), miR-995-3p (5'-TAGCACCACATGATTCGGCTT-3'), and miR-14-3p (5'-TCAGTCTTTTTCTCTCTCCTAT-3'). Fluorescent quantification was done in a LightCycler 96 Real-Time PCR System (Roche) for 50 cycles, Cts were estimated with the software provided by the manufacturer with default parameters, and ΔCts calculated using U6 spliceosomal rRNA as a normalization standard. Relative expression values in Figure 2 for unfertilized eggs were calculated with respect to the average level of bantam-3p. That is: [miR]/[bantam]=2^−ΔCt(miR)^/2^−ΔCt(bantam)^. For 2-6 hour embryos, the relative levels are calculated with respect to the levels in egg samples. Each amplification was performed in three biological replicates (independent egg/embryo collections) with two technical replicates each.

**Figure 2.**
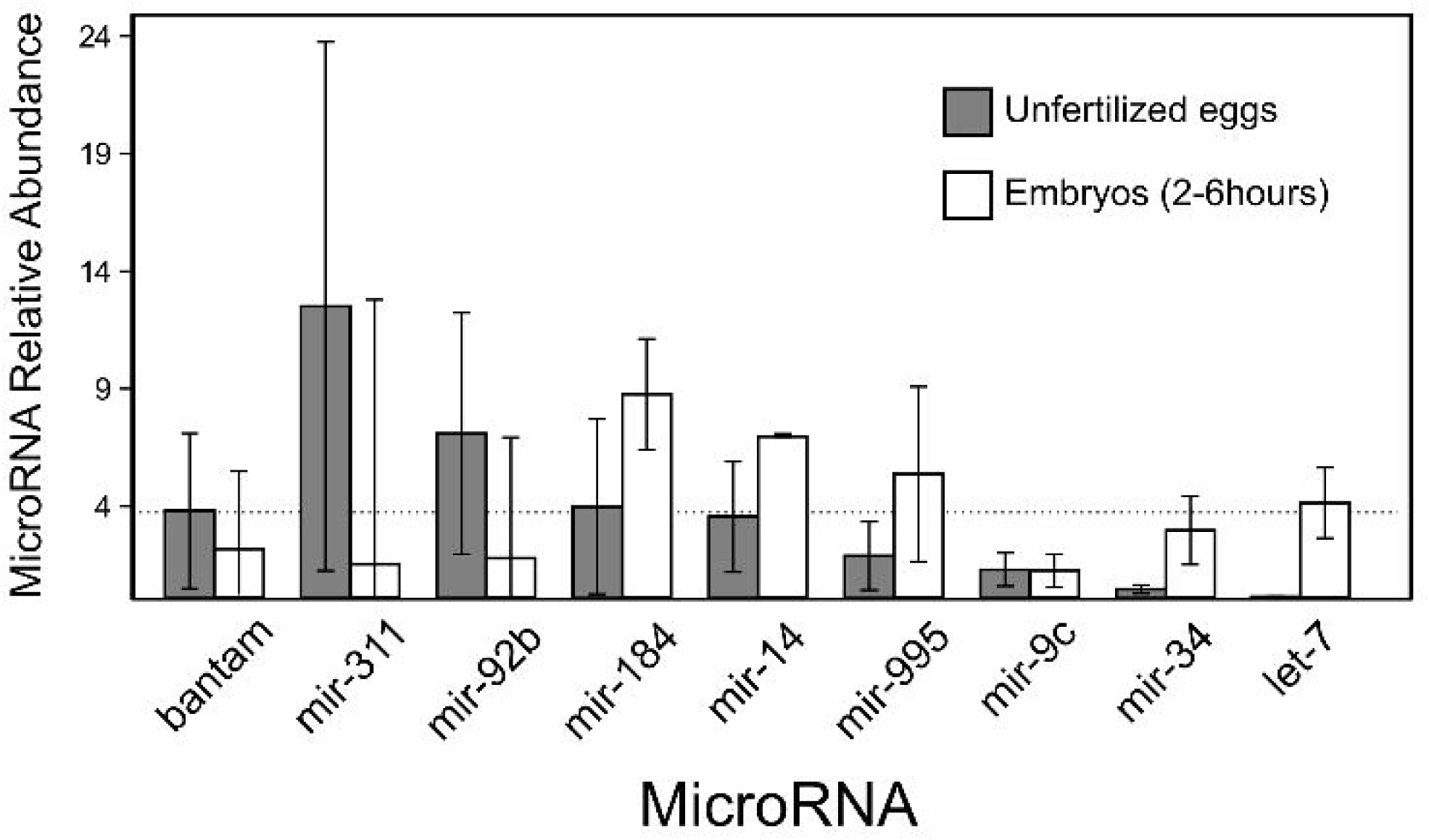
Quantification of selected maternal microRNAs in eggs and embryos. Levels of microRNA mature products in unfertilized eggs with respect with average bantam-3p levels from qPCR assays (grey boxes; see MATERIALS AND METHODS), and levels of microRNA mature products detected in 2-6 hours embryos with respect to levels in unfertilized eggs. Error bars are for three biological replicates. Dashed line indicates the levels of bantam-3p as a reference.

**MicroRNA target analysis and polymorphisms:** Target analysis was based on the presence of canonical seeds in the transcripts (Bartel 2009). Canonical seed predictions have the advantage that only primary sequence information is used, so populations models (see below) can be easily fitted. Maternally deposited gene transcripts are listed in the Berkeley Drosophila Genome Project webpage at http://insitu.fruitfly.org (Tomancak *et al.* 2007). Which transcripts are destabilized during the maternal-to-zygotic transition were identified from Tadros et al. microarray experiments (Gene Expression Omnibus accession number GSE13287), detecting probes with a >1.5 fold change in their expression level between 4-6 h embryos and oocytes (Tadros *et al.* 2007). To assess whether maternal microRNAs target transcripts that are destabilized during the MTZ transition I calculated the proportion of unstable transcripts targeted by each microRNA and compared it to the expected proportion (0.146) with a cumulative binomial test. False Discovery Rate was accounted by calculating q-values associated to the p-values (Benjamini and Hochberg 1995; Storey 2002).

For the population analyses, I first mapped the single-nucleotide polymorphisms (SNPs) from the Drosophila Genetic Reference Panel (Mackay *et al.* 2012; Huang *et al.* 2014), available at http://dgrp2.gnets.ncsu.edu/, against the 3'UTR of *Drosophila melanogaster* release 5.13 (http://flybase.org). For each microRNA I defined a target sequence (sixmer) and its 18 non-target neighbours, that is, the 18 one-nucleotide variations of the target site (Figure 3A). Every SNP that connects a target with a non-target sixmer was further considered. 3'UTRs with introns were discarded. For each polymorphic target site, the allele frequency distribution was calculated as the proportion of the target allele with respect to the total number of sampled individuals (isogenic lines). For each pair of alleles, both target and non-target sites were searched in the reference genome. This way, we also account for non-target alleles in the reference genome sequence that may be *bona fide* microRNA target sites. To study the Derived Allele Frequencies (DAF), I first mapped polymorphic target sites from *D. melanogaster* genome release 5.13 onto release 6 using the coordinate converter in FlyBase, and then finding the conserved sites in *D. sechellia* by parsing the genome sequence alignment files available at UCSC Genome Browser (ftp://hgdownload.cse.ucsc.edu/goldenPath/dm6/multiz27way; Siepel *et al.* 2005) using custom-made PERL scripts. Maternal microRNAs used in the DAF analysis mature sequences highly abundant in unfertilized eggs: bantam-3p, mir-1010-3p, mir-10-5p, mir-11-3p, mir-14-3p, mir-184-3p, mir-263a-5p, mir-276a-3p, mir-279-3p, mir-281-2-5p, mir-305-3p, mir-305-5p, mir-306-5p, mi,-313-5p, mir-318-3p, mir-31a-5p, mir-33-5p, mir-8-3p, mir-956-3p, mir-995-3p, mir-999-3p and mir-9c-5p. Non-maternal microRNAs were those not expressed in any tissue/stage according to the information available from miRBase. These microRNAs (with available SNP information) were: mir-3644-5p, mir-4941-3p, mir-4944-3p, mir-4944-5p, mir-4963-5p, mir-4967-5p, mir-4972-3p, mir-4979-5p, mir-4982-3p and mir-4985-3p.

**Figure 3.**
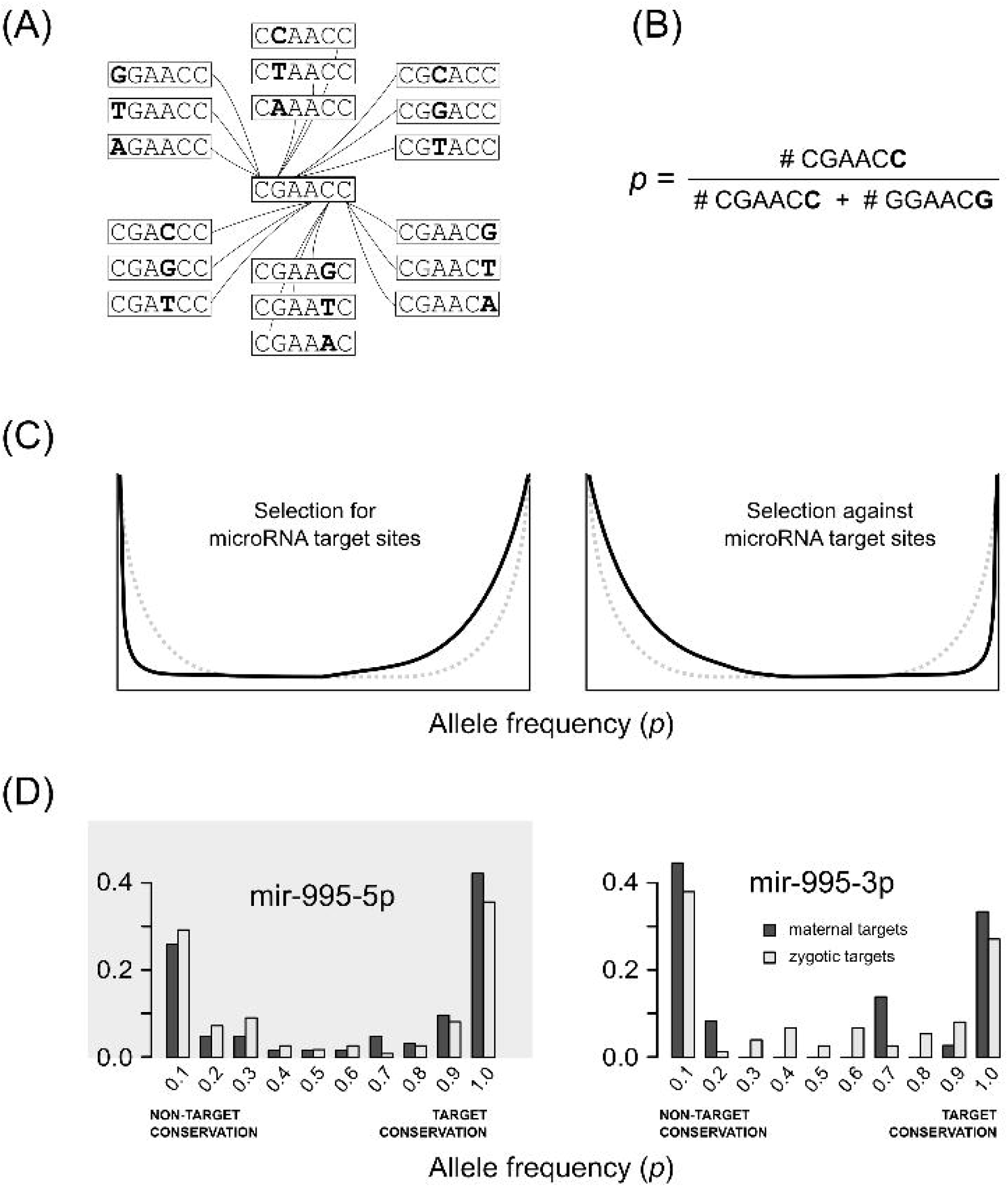
Polymorphic target sites in *Drosophila* populations. (A) Each microRNA sixmer target site has 18 one-nucleotide mutant neighbours which are themselves not target sites. (B) The allele frequency for each pair of target/non-target site is calculated as the proportion of target site alleles with respect to the total number of alleles in the pair. (C) Cartoon illustrating expected allele frequency distributions. Allele frequency distribution in a finite population is U-shaped for pairs of alleles neutral to each other (grey dashed line in both panels). If there is selection favouring target sites, distributions are expected to be shifted to the right (left panel). Conversely, if there is selection against target sites, distributions will be shifted to the left (right panel). (D) Allele frequency distribution for target sites for maternal microRNAs in maternal (dark grey) and zygotic (light grey) transcripts. Left and right panel shows the distributions for 5' and 3' arms of mir-995 respectively. Grey box in left panel indicates that mir-995-5p is virtually absent from the unfertilized egg.

## RESULTS

**Mature microRNAs are maternally deposited in the egg:** To identify maternal microRNAs in *Drosophila* I first characterized RNAs from unfertilized eggs with high-throughput sequencing (see MATERIALS AND METHODS). The most abundant microRNAs in unfertilized eggs were produced by *mir-92b*, *mir-184*, the *mir-310/mir-311/mir-312/mir-313* cluster and *bantam* genes, which accounted for over a half of the microRNA reads. Table 1 shows microRNA loci producing more than 13 reads (1‰ of the microRNA-associated reads). A full list of detected microRNAs with their read counts is in Table S1. The dataset was screened for new microRNAs as previously described (Marco *et al.* 2010; Marco, Kozomara, *et al.* 2013; Marco and Griffiths-Jones 2012), but no new microRNAs were found. This tell us that maternal microRNAs are already known in *Drosophila*.

**Table 1.** Maternal microRNAs in *Drosophila melanogaster*.

| MicroRNA transcript<sup>*</sup> | Reads per miRNA | Total reads (%)<sup>**</sup> |
| --- | --- | --- |
| mir-310/311/312/313 | 356/2012/1661/82 | 4111 (31.3) |
| mir-92a/92b | 172/2109 | 2281 (17.4) |
| mir-184 | 1377 | 1377 (10.5) |
| mir-9c/306/79/9b | 1064/154/4/132 | 1354 (10.3) |
| bantam | 1204 | 1204 (9.2) |
| mir-995 | 624 | 624 (4.8) |
| mir-14 | 411 | 411 (3.1) |
| mir-275/305 | 90/269 | 359 (2.7) |
| mir-998/11 | 5/205 | 210 (1.6) |
| mir-8 | 209 | 209 (1.6) |
| mir-2b-2/2a-1/2a-2 | 85/25/19 | 129 (1.0) |
| mir-279/996 | 61/56 | 117 (0.9) |
| mir-2b-1 | 92 | 92 (0.7) |
| mir-281-2/281-1 | 66/13 | 79 (0.6) |
| mir-4969/999 | 0/69 | 69 (0.5) |
| mir-33 | 62 | 62 (0.5) |
| mir-263a | 59 | 59 (0.4) |
| mir-10 | 28 | 28 (0.2) |
| mir-2c/13a/13b-1 | 0/0/27 | 27 (0.2) |
| mir-13b-2 | 26 | 26 (0.2) |
| mir-970 | 26 | 26 (0.2) |
| mir-1012 | 25 | 25 (0.2) |
| mir-31a | 23 | 23 (0.2) |
| mir-9a | 21 | 21 (0.2) |
| mir-309/3/286/4/5/6-1/6-2/6-3 | 0/1/15/1/1/1/1/1 | 21 (0.1) |
| mir-956 | 20 | 20 (0.2) |
| mir-276a | 18 | 18 (0.1) |
| mir-994/318 | 2/14 | 16 (0.1) |
| mir-1010 | 14 | 14 (0.1) |

\* MicroRNAs clustered in the genome (<10kb).
\*\* Percentage over total number of reads mapping to microRNAs.

In a recent report, Narry Kim and collaborators identified maternally transmitted microRNAs in *Drosophila* and demonstrated that they are targeted to degradation during maternal-to-zygotic transition (MTZ) by adenylation via Wispy (Lee *et al.* 2014). Their set of maternal microRNAs is virtually identical to the set here described. Overall, the read counts from both datasets are highly correlated (R^2^=0.62; p<0.001, Supplementary Figure S1A). The overlap for the top N-th most abundant microRNAs between both datasets is highly significant (Figure S1B). Specifically, the microRNAs here described as maternal in Table 1 (more than 13 reads) are the top 35 mature sequences, overlapping with 26 microRNAs from Lee et al's top 35 microRNAs (74.3%; p=0.00031, Figure S1B). Additionally, the read counts form this study and a recent report by Ninova *et al.* (2015), which uses the same protocol for RNA extraction and sequencing, are highly correlated (R^2^=0.86; p<0.001; Figure S1C). All these observations support the high confidence of the maternal microRNA set here described.

Figure 1 compares the relative expression of maternal microRNAs in the ovary, unfertilized eggs and early stages of development. From this comparison three types of maternal microRNAs can be distinguished. First, some maternal microRNAs are highly expressed in the ovary. A second class consists on microRNAs that are found primarily in the unfertilized egg. Third, a large proportion of maternal microRNAs is also transcribed later on during development. These groups are referred as 'high in ovary', 'high in egg' and 'high in zygote' maternal microRNAs in Figure 1. Some of these microRNAs were detected at very low levels, and whether they are bona fide maternal microRNAs may need further evidence.

To further confirm the presence of maternal microRNAs in unfertilized eggs, I validated the presence of highly abundant mature products by qPCR (see MATERIALS AND METHODS). Figure 2 shows the relative abundance of selected microRNAs (with respect to the average level of bantam-3p). Although the microRNA level varies substantially across biological replicates, the presence of 7 of the maternal microRNAs here described is validated (bantam-3p, mir-311-3p, mir-92b-3p, mir-184-3p, mir-14-3p, mir-995-3p and mir-9c-5p), although the levels of the latter two were relatively low. Furthermore, the level of mir-34-5p, which has been reported to be maternally transmitted (Soni *et al.* 2013), was very low, in agreement with this and other investigations (see Discussion). The conserved microRNA let-7-5p was used as a negative control, as it was not detected in unfertilized eggs. In the qPCR analysis, let-7-5p was not amplified in unfertilized eggs (Figure 2). I further measured the relative levels of maternal microRNAs in later stage embryos (2-6 hours). In concordance to the high-throughput sequencing analysis presented in Figure 1, bantam-3p, mir-311-3p and mir-92-3p were more abundant in the unfertilized egg than in the developing embryo. On the other hand, mir-14-3p was higher expressed in the embryo than in the egg. However, for mir-184-3p and mir-995-3p the pattern was not consistent between RNAseq and qPCR. The differences were not significant. Both mir-34-5p and let-7-5p were highly abundant in developing embryos, further supporting that they are virtually absent from the unfertilized egg and expressed from the zygotic genome at later stages during development.

**Intronic maternal microRNAs hosted in maternal protein-coding genes:** In a previous work I observed that female biased microRNAs tend to be produced from introns of female biased protein coding transcripts (Marco 2014). For instance, mir-92a is highly expressed in females, and it is encoded within the *jigr1* gene, which is maternally deposited in the egg. Here I show that mir-92a is also maternal. To further explore the relationship between maternal microRNAs and the maternal deposition of overlapping genes, I compared the expression pattern of intronic maternal microRNAs and the host protein coding gene. Table 2 lists 12 maternal microRNA clusters hosted in protein coding genes. For nine of these host genes there are in situ hybridization experiments (Tomancak *et al.* 2002, 2007), and eight of them are maternally loaded. Since 55.8% of genes in this dataset are shown to produce maternally deposited transcripts, our set of host genes is statistically enriched for maternal products (p ∼ 0.044; binomial test). There is no information from high-throughput in situ hybridization analyses for *grp*, but it is known to be present in unfertilized oocytes (Fogarty *et al.* 1997). The other two host genes have no expression information available at FlyBase. From this analysis I conclude that intronic maternal microRNAs are frequently produced from introns of maternally deposited gene transcripts.

**Table 2.**
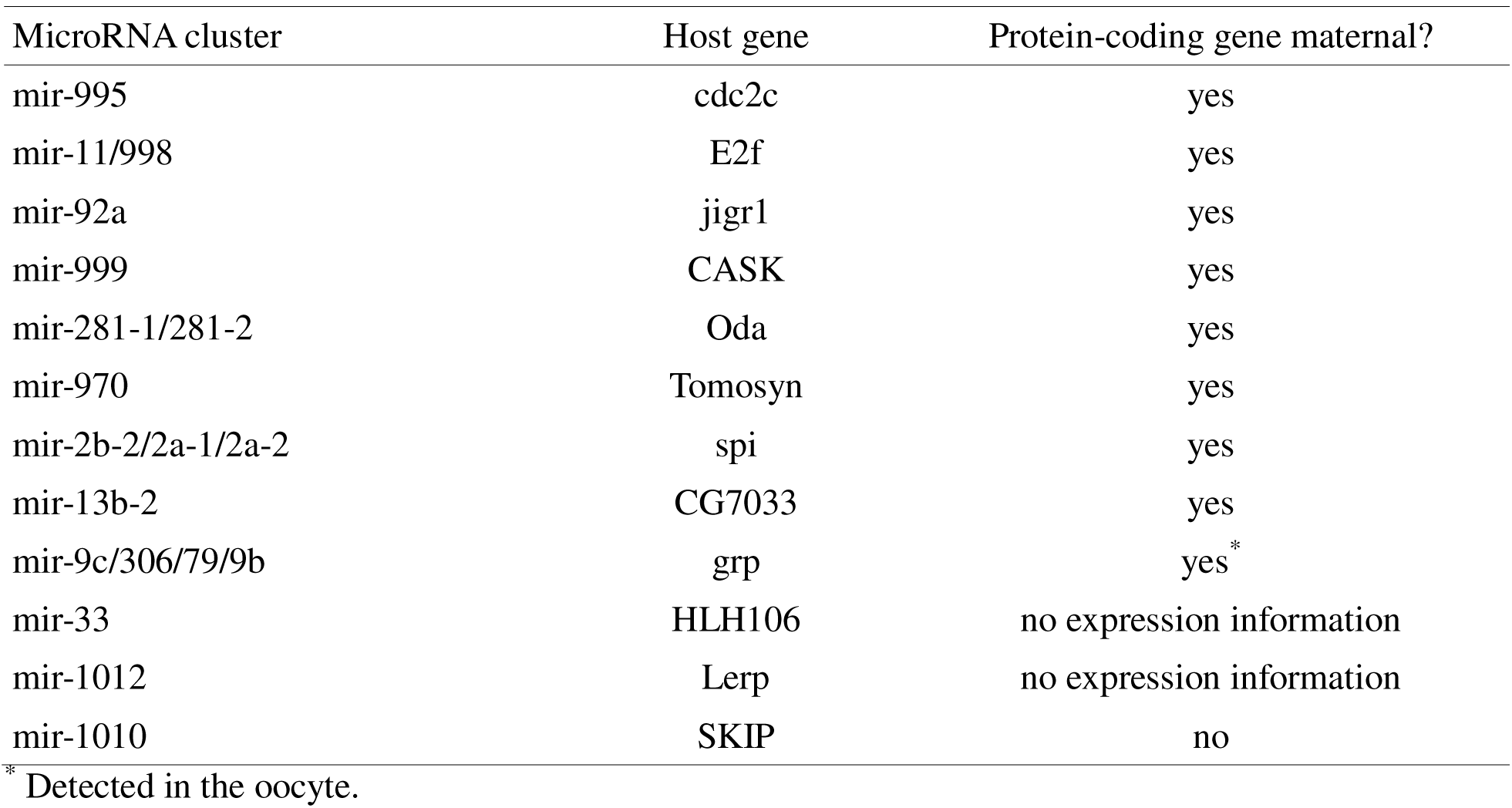
Maternal microRNA loci within protein-coding genes.

| MicroRNA cluster | Host gene | Protein-coding gene maternal? |
| --- | --- | --- |
| mir-995 | cdc2c | yes |
| mir-11/998 | E2f | yes |
| mir-92a | jigr1 | yes |
| mir-999 | CASK | yes |
| mir-281-1/281-2 | Oda | yes |
| mir-970 | Tomosyn | yes |
| mir-2b-2/2a-1/2a-2 | spi | yes |
| mir-13b-2 | CG7033 | yes |
| mir-9c/306/79/9b | grp | yes* |
| mir-33 | HLH106 | no expression information |
| mir-1012 | Lerp | no expression information |
| mir-1010 | SKIP | no |

\* Detected in the oocyte.

**Maternal microRNAs in the Maternal-to-Zygotic Transition:** As shown in Figure 1, a significant fraction of maternal microRNAs have a lower expression when zygotic transcription starts. One possibility is that some of these maternal microRNAs have a role in destabilizing maternal transcripts during the MZT. A similar role has been described for early expressed zygotic microRNAs in *Drosophila* (Bushati *et al.* 2008) and other species such as zebrafish (Giraldez *et al.* 2006). I predicted target sites for each maternal microRNAs in stable and unstable maternal transcripts during MZT (Tadros *et al.* 2007). Table 3 shows maternal microRNAs targeting more unstable maternal transcripts than expected by chance (FDR < 10%). Two of the microRNAs, mir-283 and mir-277, were detected at very low levels in unfertilized eggs (Table S1) and have a higher expression level later on during embryonic development (Figure 1). It is possible that these microRNAs contribute to the destabilization of maternal transcripts, but probably as zygotic microRNAs. Another set of microRNAs which may contribute to transcript clearance during MZT are the mir-310 and mir-92 families. They both share the same seed sequence (which determines the targeted transcripts). These are also zygotic microRNAs expressed very early during development. Last, the microRNA-9 family also target unstable maternal transcripts. Members of the mir-9 family, particularly mir-9c, are particularly abundant in unfertilized eggs but lower expressed in early embryos (Figure 1). This indicates that mir-9 may be the first case of a maternal microRNA contributing to the degradation of maternal transcripts during MZT. In summary, some maternally deposited microRNAs have a potential role in destabilizing maternal transcripts.

**Table 3.**
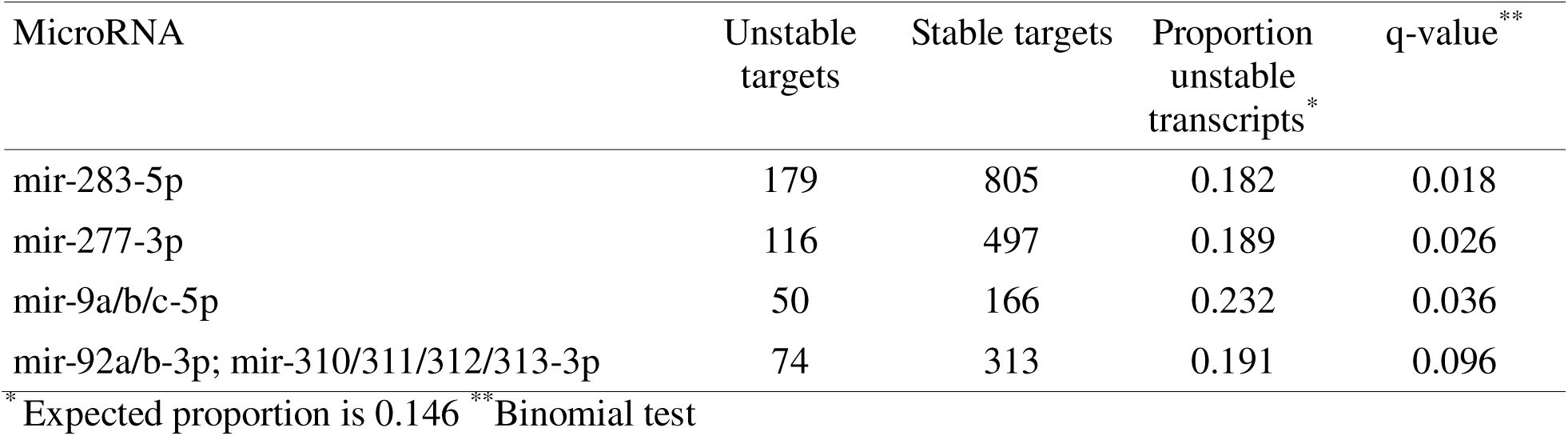
Maternal microRNAs targeting unstable transcripts during maternal-to-zygotic transition

**Maternal protein-coding transcripts are selected against target sites for maternal microRNAs:** If maternal microRNAs have a functional impact on maternal transcripts, these transcripts should have a different target repertoire compared to zygotic transcripts. I estimated how many maternal and zygotic transcripts are targeted by maternal microRNAs. Overall, 73% of maternal transcripts and 63% of zygotic transcripts have canonical seed target sites for maternal microRNAs. However, for transcripts from genes with a recent evolutionary origin, that is, that they originated in the *Drosophila melanogaster* lineage, maternal transcripts were less likely to be targeted by maternal microRNAs than zygotic transcripts: 50.7±0.5% of maternal transcripts have canonical target sites for maternal microRNAs, whilst this percentage is 52.6±0.4% for zygotic transcripts (p∼0.004; t-test). Although the difference is small, the observation that evolutionarily young maternal genes have a relatively lower proportion of targets for maternal microRNAs than zygotic genes suggest purifying selection against microRNA targets. In other words, if there was no selection against microRNA targets, we would expect a similar proportion of target sites between maternal and zygotic transcripts.

To test whether there is selection against maternal microRNA target sites we should evaluate population data. To do so, I first constructed a model of microRNA target mutation as follows (see Figure 3A): 1) a target site is defined as any 6 nucleotide sequence (sixmer) in a 3'UTR complementary to the seed region (Bartel 2009) of a microRNA; 2) any target site has 18 mutant neighbours, which are one nucleotide mutation apart from the canonical target, and are not themselves targets; 3) only polymorphic sites in which one of the alleles is a target site and the other a non-target are further considered in this analysis. Allele frequency is here defined as the proportion of the target allele (*p* in Figure 3B). For instance, an allele frequency of 0.8 means that 80% of the sampled individuals have the target site at a given position and 20% have a non-target mutant neighbour. Conversely, an allele frequency of 0.3 will indicate that the non-target neighbour is more frequent (70%) than the target allele (30%). Population genetics theory (Crow and Kimura 1970; Nei 1975) predicts that, in a finite population, two alleles neutral to each other will have a symmetric U-shaped distribution, that is, most individuals will be homozygous for one of the alleles. However, if there is a selective pressure to conserve a target site, the distribution will be shifted to the right. On the other hand, if selection is against a target site allele, the distribution will be shifted to the left (see Figure 3C). A symmetric U-shape distribution is not expected if other evolutionary forces are in place (for instance, mutation bias, or background selection produced by purifying selection on neighbouring sites). Hence, in order to estimate the selective pressure for, or against, a microRNA target site in maternal transcripts, we need an empirical expected distribution of allele frequencies. Therefore, I calculated the allele frequency at target sites in zygotic transcripts, in which maternal microRNAs have no (or little) influence. By comparing the allele frequency distribution of target sites between maternal and zygotic transcripts we can estimate the relative selective pressure on microRNA target sites in maternal with respect to zygotic transcripts.

Figure 3D shows the case for *mir-995* microRNA products. One of them, mir-995-3p is abundant in unfertilized eggs whilst the alternate arm, mir-995-5p, is virtually absent in eggs. The allele frequency distribution in maternal transcripts is shifted to the left with respect to zygotic transcripts in mir-995-3p. That is not the case for mir-995-5p. In other words, there is a preference for alleles that are non-targets of maternal mir-995-3p, but not for the non-maternal mir-995-5p. Both arms of mir-305 are present at high levels in unfertilized eggs. Figure 4A shows the allele frequency distribution for their targets, and both arms show evidence of selective pressure against maternal microRNA target sites. As a counterexample, Figure 4B shows the allele frequency distribution of a microRNA for which none of the arms was detected in unfertilized eggs: mir-4986. Consistently, none of the microRNA products showed evidence of selection against target sites.

**Figure 4.**
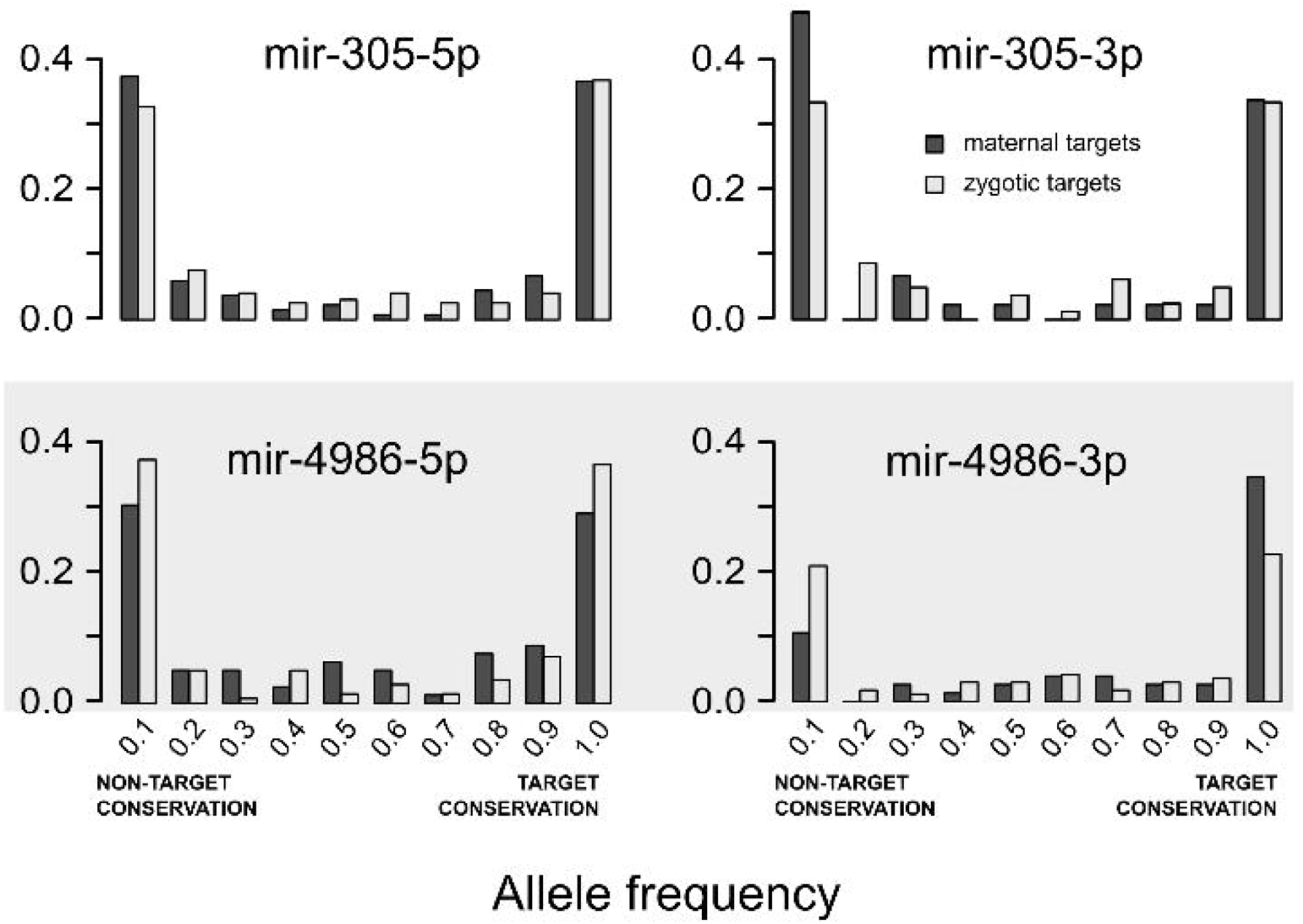
Allele frequency distribution for target sites of maternal microRNAs. Distribution of targets for maternal and zygotic transcripts for mir-305 (both arms are highly present in the egg) and mir-4986 (grey box, neither of the arms was detected in the egg).

To explore whether this pattern is a general feature of maternal microRNAs I defined 'target avoidance' as the log2 ratio of the number of target sites with an allele frequency smaller than 0.1 (that is, the frequency of sites where >90% of alleles are the non-target sequence) between maternal and zygotic transcripts. In this context, positive values indicate that targets for a specific microRNA tend to be 'avoided' by maternal transcripts, that is, there is selection against target sites for maternal microRNAs in maternal transcripts. Figure 5A is a bar plot of target avoidance values for different levels of microRNA abundance in the egg. Maternally deposited coding transcripts tend to avoid some target sites for highly abundant maternal microRNAs (with respect to zygotic transcripts). Differences were statistically significant (Figure 5A). In a similar manner I defined 'target conservation' as the log2 ratio of the number of target sites with allele frequency greater than 0.9 between maternal and zygotic transcripts. A positive value indicates that target-sites are preferentially conserved in maternal transcripts. Figure 5B shows these values for different microRNA abundances. Overall, maternal transcripts conserve some target sites, but there is not a distinctive profile between maternal and non-maternal microRNAs (Figure 5B).

**Figure 5.**
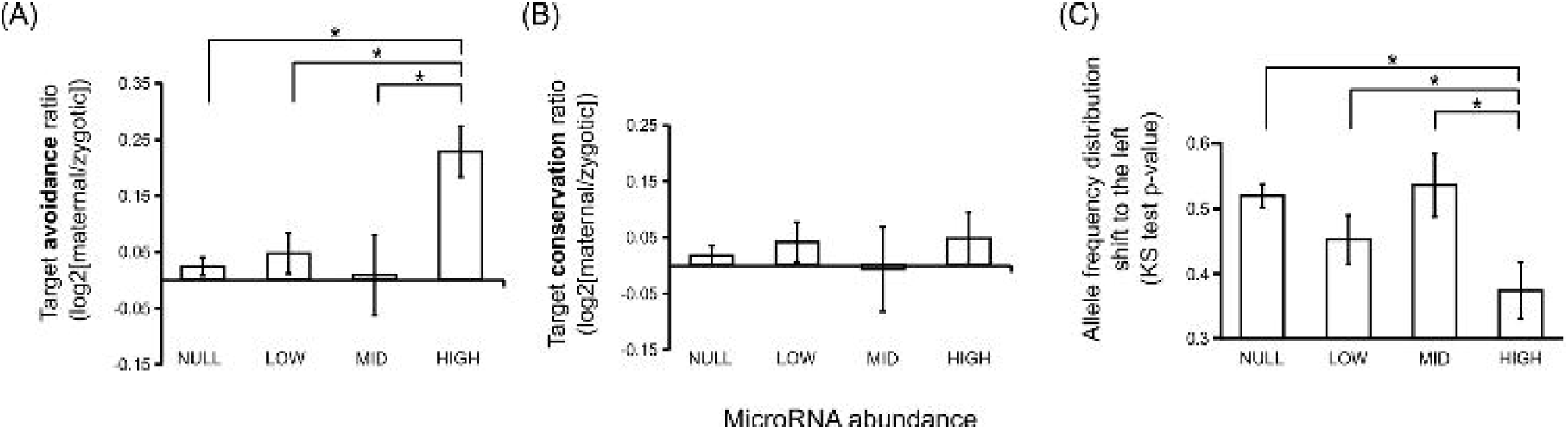
Maternal microRNA target avoidance. (A) Target avoidance (see main text for details) for microRNAs with differences abundances in the unfertilized egg (NULL – not detected, LOW – less than 0.1% of the set, MID – between 0.1% and 1%, HIGH – more than 1%). Error bars represent the Standard Error of the Mean. Asterisks show statistically significant differences (p<0.01) for t-test with unequal variances. (B) Target conservation for microRNAs with differences abundances in the unfertilized egg. (C) Distribution shifting to the left of allele frequency distribution in maternal transcripts with respect to zygotic transcripts.

We can also compare the whole allele frequency distribution and evaluate whether the distribution for maternal transcripts is shifted to the left with respect to that of zygotic transcripts (as suggested in Figure 3C, right panel). To do so I performed a one-tail Kolmogorov-Smirnov (KS) test for each pair of allele frequency distributions, and evaluated whether the distribution of maternal alleles was shifted to the left compared to the zygotic allele distribution: that is, whether there is a preference for the non-target allele in maternal transcripts. The lower the p-value, the larger the shift to the left. This measure is not independent from that in Figure 5A, and I use it here as an alternative method to evaluate target avoidance. Figure 5C plots the KS p-value, and shows that these are significantly lower for microRNAs that are abundant in unfertilized eggs. The allele frequency distributions of maternal microRNA target sites in maternal transcripts are, therefore, biased towards the non-target allele.

The comparison of two allele frequency distributions has been very useful to detect selection and/or mutational biases (Galtier *et al.* 2006; Eyre-Walker *et al.* 2006). Nevertheless, it is difficult to infer directionality in the evolutionary process as we do not know which one was the ancestral allele. One way to circumvent this issue is to compute the Derived Allele Frequency (DAF) distribution, which is the allele frequency distribution of alleles that were not ancestral. The comparison of DAF distributions has been of much use to infer selection in genes (Yngvadottir *et al.* 2009) and in regulatory sites (Sethupathy *et al.* 2008). I computed the DAF distribution for non-target sites to explore signatures of selection against target sites in maternal transcripts. First, I catalogued, among all target/non-target pair of alleles analysed in this study, those sites that were conserved in *Drosophila sechellia*, which diverged from the *D. melanogaster* lineage about 2 million years ago. Then, I selected those polymorphic sites that are non-target sites in *D. sechellia* and, assuming that that was the ancestral state, I plotted the frequency distribution of the target alleles in *D. melanogaster*. As maternal microRNAs I selected highly abundant mature sequences. As non-maternal I selected microRNAs that were not present in the egg, but also low or no expressed in other tissues (see MATERIALS AND METHODS). Figure 6 compares the DAF for ancestral non-target sites in maternal transcripts between maternal and non-maternal microRNAs. In agreement with the previous analyses, the derived allele frequencies are smaller for maternal than for non-maternal microRNA targets. The difference between the two distributions was significant (p<0.0001; Kolmogorov-Smirnov test). Also, there was an excess of singletons in maternal (67 out of 236) compared to non-maternal (44 out of 223) microRNA target sites (p=0.0304; Chi-square test). The whole dataset is available in File S1. That indicates that selection may favour the derived allele, that is, the non-target allele. In conclusion, different analyses suggest purifying selection against maternal microRNA target sites in maternal transcripts.

**Figure 6.**
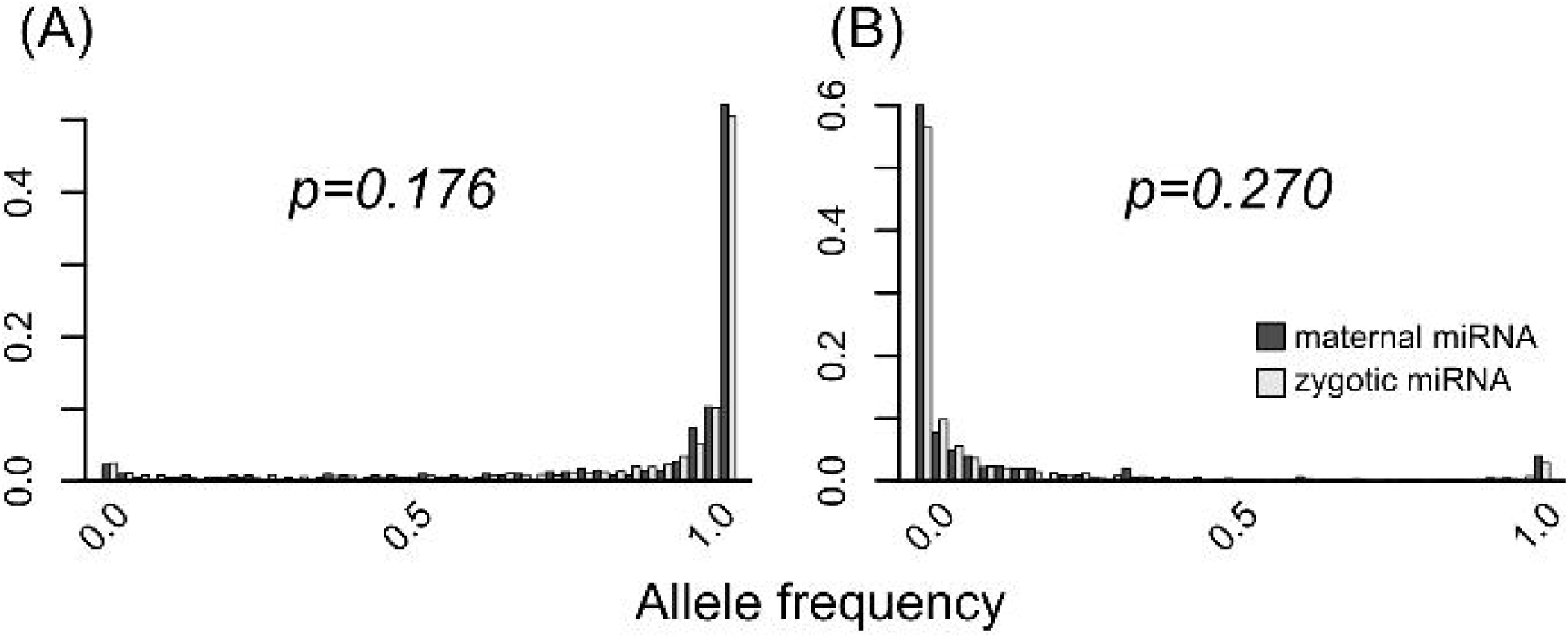
Derived Allele Frequency distribution of microRNA target sites. Allele frequency distribution of SNPs that are microRNA targets whose predicted ancestral state was a non-target site. The frequency distribution for maternal microRNA target sites is plotted in dark grey boxes, and the distribution for non-maternal microRNA target sites in light grey boxes.

## DISCUSSION

This study characterizes microRNA products from *Drosophila* unfertilized eggs. I validated seven of these microRNAs by qPCR. The presence of microRNAs in unfertilized oocytes have been described in mice (Tang *et al.* 2007). However, it has been shown that microRNA activity is suppressed in mice oocytes, indicating that maternally deposited microRNAs may not have a defined function in this species (Ma *et al.* 2010; Suh *et al.* 2010). Here I show evidence for *Drosophila* maternal microRNA activity as they have an impact in the evolution of potential target sites in maternal microRNAs (Figures 3, 4, 5 and 6).

One of the most abundant maternal microRNAs, mir-184, has been already described in freshly laid *Drosophila* eggs (Iovino *et al.* 2009). Also, the *mir-184* gene has an important role during oocyte development as well as in early development (Iovino *et al.* 2009). Another maternal microRNA gene, *mir-14*, seems to be involved in transcriptional silencing of transposable elements in the germline (Mugat *et al.* 2015). On the other hand, the conserved microRNA mir-34 has been also described as a maternal microRNA (Soni *et al.* 2013) but it has only one read copy in our dataset, and it has not been detected in two other independent high-throughput screens (Lee *et al.* 2014; Ninova *et al.* 2015). The level of mir-34-5p was also very low in specific qPCR assays (Figure 2). All these suggest that either mir-34 is a very low copy maternal microRNA, or that it is rapidly degraded after egg deposition/activation.

Another maternal microRNA gene, *mir-9c*, is necessary to regulate the number of germ cells (Kugler *et al.* 2013). Indeed, is the maternal loss of *mir-9c* what produces this phenotype (Kugler *et al.* 2013). This microRNA is hosted within a maternally deposited gene, *grapes* (Table 2). Here I show that mir-9c-5p targets more unstable transcript during the MZT than expected by chance (Table 3), which indicates that mir-9c-5p may have a role during maternal transcript clearance during the initial steps of development. A similar role has been described for zygotically transcribed microRNAs (Bushati *et al.* 2008).

Other maternally deposited microRNAs derive from the *mir-310/mir-313* cluster. This cluster is highly conserved in the *Drosophila* lineage (Marco, Ninova, *et al.* 2013), although it may have originated in insects (Ninova *et al.* 2014), and is evolutionarily related with the (also maternal) *mir-92a/mir-92b* cluster (Lu *et al.* 2008; Ninova *et al.* 2014). Mature products from the orthologous *mir-310/311/312/313* and *mir-92a/92b* clusters in *Drosophila virilis* has been detected at high levels during the first two hours of development, suggesting that this cluster is also maternally deposited in this species (Supplementary Table 2 in (Ninova *et al.* 2014)). Interestingly, some maternal microRNAs have other functions later on during development. MicroRNAs from the *mir-310/311/312/313* cluster are known to be involved in male gonad development (Pancratov *et al.* 2013). Recently, Ranz and collaborators found that mir-310/mir-313 microRNAs show male biased expression pattern at the onset of metamorphosis (Yeh *et al.* 2014). On the other hand, *mir-92a* is expressed in the adult, and it is involved in leg morphology (Arif *et al.* 2013). Some other maternal microRNAs have roles unrelated with embryonic development such as *mir-14*, which regulates insulin production (Xu *et al.* 2003); *mir-279*, involved in the circadian clock (Luo and Sehgal 2012 p. 279); or *mir-8*, associated to abdominal pigmentation (Kennell *et al.* 2012), to name but a few cases. Altogether, these examples show that maternal microRNAs frequently have other functions at different developmental stages and/or tissues.

MicroRNA target avoidance has been observed in *Drosophila* (Stark *et al.* 2005), as well as in mice (Farh *et al.* 2005) and humans (Sood *et al.* 2006; Chen and Rajewsky 2006). Here I detect a similar pattern in *Drosophila* eggs, in which maternal transcripts tend to avoid target sites for maternal microRNAs. Alternatively, a lower number of target sites in maternal transcripts may be explained as an early degradation of transcripts with conserved target sites, and therefore not detected in early embryos. However, in *Drosophila*, microRNA-mediated transcript degradation happens a few hours after microRNA-mediated repression (Djuranovic *et al.* 2012). Maternal transcripts are detected from 0-2 hour old embryos, and they are unlikely to have had microRNA-mediated transcript degradation. The microRNA genes studied in that paper were: *mir-9b*, *mir-279* and *bantam*, all of which were detected in this study as maternal.

If microRNAs are likely to have a function in maternal transcripts, why we observe selection against target sites? I suggest the following explanation. A microRNA that is maternally deposited and targets several maternal microRNAs may have a function, for instance, induce the programmed degradation of maternal transcripts during MZT. However, there are hundreds of other maternal transcripts that should not be targeted. This situation creates a conflict in which functional interactions must be conserved, but new interactions that potentially impair existing regulatory networks should be avoided. In this context, most maternal transcripts will be selected against target sites for maternal microRNAs. It is likely that this conflict also happens in other tissues and species, and probably will also affect transcription factor mediated regulation. How much selection against regulatory sites affects genome evolution is not yet known, and more studies need to be done.

The main advantage of working with microRNAs to study evolution at the population level is that we can predict the impact of single point mutations in both the microRNAs and their targets. This is not yet possible with other gene regulators, such as transcription factors. I introduce a simple mutation model to study target/non-target allele pairs, and propose that comparing the allele frequencies at target sites between two groups of targeted genes can be use to infer selective pressures on microRNA target sites. The use of population genetics to study the evolutionary dynamics of microRNA target sites is still an underdeveloped research area. Despite the limitations of the model here introduced, it has been proved to be useful to detect selection at microRNA target sites. I anticipate that more accurate models and the analyses of bigger sets of microRNA target sites will shed light on how microRNA function diversify and, more generally, how gene regulation evolves.

Overall, this paper describes three features of maternally transmitted microRNAs: 1) they are often produced from introns of maternally deposited transcripts; 2) they can be zygotically transcribed and have other functions during development; 3) maternal transcripts tend to avoid target sites for maternal microRNAs. Additionally, I suggest that mir-9c may be involved in maternal transcript clearance during MZT. These observations indicate that some maternal microRNAs may have a function but are potentially damaging to the normal function of other maternal genes. Therefore, selective pressures may prevent maternal transcripts to be targeted by maternal microRNAs.

## ACKNOWLEDGEMENTS

I thank Maria Ninova and Fran Bonath for sharing their expertise on small RNA library preparations, and to Matt Ronshaugen for helpful advice on fly genetics. Greg Brooke, Elena Klenova and Adele Angel hosted me in their lab. I am also very grateful to Maria Ninova and Sam Griffiths-Jones form critical reading of the manuscript, and to my colleagues at the Junior European Drosophila Investigators (JEDI) network for useful feedback. John Kim, Stephen Wright and two anonymous reviewers made critical and important comments on earlier versions of this manuscript. The MiSeq sequencing was funded by the Career Developmental Award (University of Manchester). Real-time qPCR assays and publication costs were covered by the University of Essex.

## Reference

Aboobaker, A. A., P. Tomancak, N. Patel, G. M. Rubin, and E. C. Lai, 2005 Drosophila microRNAs exhibit diverse spatial expression patterns during embryonic development. Proc. Natl. Acad. Sci. U. S. A. 102: 18017–22.

Aravin, A. A., M. Lagos-Quintana, A. Yalcin, M. Zavolan, D. Marks et al., 2003 The small RNA profile during Drosophila melanogaster development. Dev. Cell 5: 337–350.

Arif, S., S. Murat, I. Almudi, M. D. S. Nunes, D. Bortolamiol-Becet et al., 2013 Evolution of mir-92a Underlies Natural Morphological Variation in Drosophila melanogaster. Curr. Biol. 23: 523–528.

Bartel, D. P., 2009 MicroRNAs: target recognition and regulatory functions. Cell 136: 215–233.

Benjamini, Y., and Y. Hochberg, 1995 Controlling the False Discovery Rate: A Practical and Powerful Approach to Multiple Testing. J. R. Stat. Soc. Ser. B Methodol. 57: 289–300.

Bushati, N., A. Stark, J. Brennecke, and S. M. Cohen, 2008 Temporal reciprocity of miRNAs and their targets during the maternal-to-zygotic transition in Drosophila. Curr. Biol. CB 18: 501–506.

Chen, P. Y., H. Manninga, K. Slanchev, M. Chien, J. J. Russo et al., 2005 The developmental miRNA profiles of zebrafish as determined by small RNA cloning. Genes Dev. 19: 1288–1293.

Chen, K., and N. Rajewsky, 2006 Natural selection on human microRNA binding sites inferred from SNP data. Nat. Genet. 38: 1452–1456.

Crow, J. F., and M. Kimura, 1970 An Introduction to Population Genetics Theory. Harper & Row.

Czech, B., C. D. Malone, R. Zhou, A. Stark, C. Schlingeheyde et al., 2008 An endogenous small interfering RNA pathway in Drosophila. Nature 453: 798–802.

Djuranovic, S., A. Nahvi, and R. Green, 2012 miRNA-Mediated Gene Silencing by Translational Repression Followed by mRNA Deadenylation and Decay. Science 336: 237–240.

Enright, A., B. John, U. Gaul, T. Tuschl, C. Sander et al., 2003 MicroRNA targets in Drosophila. Genome Biol. 5: R1.

Eyre-Walker, A., M. Woolfit, and T. Phelps, 2006 The Distribution of Fitness Effects of New Deleterious Amino Acid Mutations in Humans. Genetics 173: 891–900.

Farh, K. K.-H., A. Grimson, C. Jan, B. P. Lewis, W. K. Johnston et al., 2005 The widespread impact of mammalian MicroRNAs on mRNA repression and evolution. Science 310: 1817–1821.

Fogarty, P., S. D. Campbell, R. Abu-Shumays, B. de Saint Phalle, K. R. Yu et al., 1997 The Drosophila grapes gene is related to checkpoint gene chk-1 rad27 and is required for late syncytial division fidelity. Curr. Biol. 7: 418–426.

Galtier, N., E. Bazin, and N. Bierne, 2006 GC-biased segregation of noncoding polymorphisms in Drosophila. Genetics 172: 221–228.

Giraldez, A. J., Y. Mishima, J. Rihel, R. J. Grocock, S. Van Dongen et al., 2006 Zebrafish MiR-430 promotes deadenylation and clearance of maternal mRNAs. Science 312: 75–79.

Huang, W., A. Massouras, Y. Inoue, J. Peiffer, M. Ràmia et al., 2014 Natural variation in genome architecture among 205 Drosophila melanogaster Genetic Reference Panel lines. Genome Res. 24: 1193–1208.

Iovino, N., A. Pane, and U. Gaul, 2009 miR-184 Has Multiple Roles in Drosophila Female Germline Development. Dev. Cell 17: 123–133.

Kennell, J. A., K. M. Cadigan, I. Shakhmantsir, and E. J. Waldron, 2012 The microRNA miR-8 is a positive regulator of pigmentation and eclosion in Drosophila. Dev. Dyn. Off. Publ. Am. Assoc. Anat. 241: 161–168.

Kozomara, A., and S. Griffiths-Jones, 2014 miRBase: annotating high confidence microRNAs using deep sequencing data. Nucleic Acids Res. 42: D68–73.

Kugler, J.-M., Y.-W. Chen, R. Weng, and S. M. Cohen, 2013 Maternal Loss of miRNAs Leads to Increased Variance in Primordial Germ Cell Numbers in Drosophila melanogaster. G3 GenesGenomesGenetics 3: 1573–1576.

Lai, E. C., P. Tomancak, R. W. Williams, and G. M. Rubin, 2003 Computational identification of Drosophila microRNA genes. Genome Biol. 4: R42.

Langmead, B., C. Trapnell, M. Pop, and S. L. Salzberg, 2009 Ultrafast and memory-efficient alignment of short DNA sequences to the human genome. Genome Biol. 10: R25.

Lawrence, P. A., 1992 The Making of a Fly: The Genetics of Animal Design. Blackwell Scientific.

Lee, M., Y. Choi, K. Kim, H. Jin, J. Lim et al., 2014 Adenylation of Maternally Inherited MicroRNAs by Wispy. Mol. Cell 56: 696–707.

Lee, Y. S., K. Nakahara, J. W. Pham, K. Kim, Z. He et al., 2004 Distinct Roles for Drosophila Dicer-1 and Dicer-2 in the siRNA/miRNA Silencing Pathways. Cell 117: 69–81.

Lu, J., Y. Fu, S. Kumar, Y. Shen, K. Zeng et al., 2008 Adaptive evolution of newly emerged micro-RNA genes in Drosophila. Mol. Biol. Evol. 25: 929–938.

Luo, W., and A. Sehgal, 2012 microRNA-279 acts through the JAK/STAT pathway to regulate circadian behavioral output in Drosophila. Cell 148: 765–779.

Mackay, T. F. C., S. Richards, E. A. Stone, A. Barbadilla, J. F. Ayroles et al., 2012 The Drosophila melanogaster Genetic Reference Panel. Nature 482: 173–178.

Ma, J., M. Flemr, P. Stein, P. Berninger, R. Malik et al., 2010 MicroRNA activity is suppressed in mouse oocytes. Curr. Biol. CB 20: 265–270.

Marco, A., 2012 Regulatory RNAs in the light of Drosophila genomics. Brief. Funct. Genomics 11: 356–365.

Marco, A., 2014 Sex-biased expression of microRNAs in Drosophila melanogaster. Open Biol. 4: 140024.

Marco, A., and S. Griffiths-Jones, 2012 Detection of microRNAs in color space. Bioinformatics 28: 318–323.

Marco, A., J. H. L. Hui, M. Ronshaugen, and S. Griffiths-Jones, 2010 Functional shifts in insect microRNA evolution. Genome Biol. Evol. 2: 686–696.

Marco, A., A. Kozomara, J. H. L. Hui, A. M. Emery, D. Rollinson et al., 2013 Sex-Biased Expression of MicroRNAs in Schistosoma mansoni. PLoS Negl Trop Dis 7: e2402.

Marco, A., M. Ninova, M. Ronshaugen, and S. Griffiths-Jones, 2013 Clusters of microRNAs emerge by new hairpins in existing transcripts. Nucleic Acids Res. 41: 7745–7752.

Mugat, B., A. Akkouche, V. Serrano, C. Armenise, B. Li et al., 2015 MicroRNA-Dependent Transcriptional Silencing of Transposable Elements in Drosophila Follicle Cells. PLoS Genet 11: e1005194.

Nakahara, K., K. Kim, C. Sciulli, S. R. Dowd, J. S. Minden et al., 2005 Targets of microRNA regulation in the Drosophila oocyte proteome. Proc. Natl. Acad. Sci. U. S. A. 102: 12023–12028.

Nei, M., 1975 Molecular Population Genetics and Evolution. North-Holland.

Ninova, M., M. Ronshaugen, and S. Griffiths-Jones, 2014 Fast-evolving microRNAs are highly expressed in the early embryo of Drosophila virilis. RNA 20: 360–372.

Ninova, M., M. Ronshaugen, and S. Griffiths-Jones, 2015 Tribolium castaneum as a model for microRNA evolution, expression and function during short germband development. bioRxiv 018424.

Pancratov, R., F. Peng, P. Smibert, J.-S. Yang, E. R. Olson et al., 2013 The miR-310/13 cluster antagonizes β-catenin function in the regulation of germ and somatic cell differentiation in the Drosophila testis. Dev. Camb. Engl. 140: 2904–2916.

R Development Core Team, 2004 R: A language and environment for statistical computing.

Robinson, S. W., P. Herzyk, J. A. T. Dow, and D. P. Leader, 2013 FlyAtlas: database of gene expression in the tissues of Drosophila melanogaster. Nucleic Acids Res. 41: D744–D750.

Ruby, J. G., C. Jan, C. Player, M. J. Axtell, W. Lee et al., 2006 Large-Scale Sequencing Reveals 21U-RNAs and Additional MicroRNAs and Endogenous siRNAs in C. elegans. Cell 127: 1193–1207.

Ruby, J. G., A. Stark, W. K. Johnston, M. Kellis, D. P. Bartel et al., 2007 Evolution, biogenesis, expression, and target predictions of a substantially expanded set of Drosophila microRNAs. Genome Res. 17: 1850–1864.

Seitz, H., M. Ghildiyal, and P. D. Zamore, 2008 Argonaute Loading Improves the 5' Precision of Both MicroRNAs and Their miRNA* Strands in Flies. Curr. Biol. 18: 147–151.

Sethupathy, P., H. Giang, J. B. Plotkin, and S. Hannenhalli, 2008 Genome-wide analysis of natural selection on human cis-elements. PLoS ONE 3: e3137.

Siepel, A., G. Bejerano, J. S. Pedersen, A. S. Hinrichs, M. Hou et al., 2005 Evolutionarily conserved elements in vertebrate, insect, worm, and yeast genomes. Genome Res. 15: 1034–1050.

Soni, K., A. Choudhary, A. Patowary, A. R. Singh, S. Bhatia et al., 2013 miR-34 is maternally inherited in Drosophila melanogaster and Danio rerio. Nucleic Acids Res. 41: 4470–4480.

Sood, P., A. Krek, M. Zavolan, G. Macino, and N. Rajewsky, 2006 Cell-type-specific signatures of microRNAs on target mRNA expression. Proc. Natl. Acad. Sci. U. S. A. 103: 2746–2751.

Stark, A., J. Brennecke, N. Bushati, R. B. Russell, and S. M. Cohen, 2005 Animal MicroRNAs confer robustness to gene expression and have a significant impact on 3’UTR evolution. Cell 123: 1133–1146.

Storey, J. D., 2002 A direct approach to false discovery rates. J. R. Stat. Soc. Ser. B Stat. Methodol. 64: 479–498.

Suh, N., L. Baehner, F. Moltzahn, C. Melton, A. Shenoy et al., 2010 MicroRNA function is globally suppressed in mouse oocytes and early embryos. Curr. Biol. CB 20: 271–277.

Tadros, W., A. L. Goldman, T. Babak, F. Menzies, L. Vardy et al., 2007 SMAUG Is a Major Regulator of Maternal mRNA Destabilization in Drosophila and Its Translation Is Activated by the PAN GU Kinase. Dev. Cell 12: 143–155.

Tang, F., M. Kaneda, D. O’Carroll, P. Hajkova, S. C. Barton et al., 2007 Maternal microRNAs are essential for mouse zygotic development. Genes Dev. 21: 644–648.

Tesfaye, D., D. Worku, F. Rings, C. Phatsara, E. Tholen et al., 2009 Identification and expression profiling of microRNAs during bovine oocyte maturation using heterologous approach. Mol. Reprod. Dev. 76: 665–677.

Tomancak, P., A. Beaton, R. Weiszmann, E. Kwan, S. Shu et al., 2002 Systematic determination of patterns of gene expression during Drosophila embryogenesis. Genome Biol. 3: RESEARCH0088.

Tomancak, P., B. P. Berman, A. Beaton, R. Weiszmann, E. Kwan et al., 2007 Global analysis of patterns of gene expression during Drosophila embryogenesis. Genome Biol. 8: R145.

Tsien, H. C., and J. M. Wattiaux, 1971 Effect of Maternal Age on DNA and RNA Content of Drosophila Eggs. Nature 230: 147–148.

Votruba, S. M., 2009 MicroRNAS in the Drosophila Egg and Early Embryo. University of Toronto.

Watanabe, T., A. Takeda, K. Mise, T. Okuno, T. Suzuki et al., 2005 Stage-specific expression of microRNAs during Xenopus development. FEBS Lett. 579: 318–324.

Xu, P., S. Y. Vernooy, M. Guo, and B. A. Hay, 2003 The Drosophila microRNA Mir-14 suppresses cell death and is required for normal fat metabolism. Curr. Biol. CB 13: 790–795.

Yeh, S.-D., M. von Grotthuss, K. A. Gandasetiawan, S. Jayasekera, X.-Q. Xia et al., 2014 Functional Divergence of the miRNA Transcriptome at the Onset of Drosophila Metamorphosis. Mol. Biol. Evol. 31: 2557–2572.

Yngvadottir, B., Y. Xue, S. Searle, S. Hunt, M. Delgado et al., 2009 A Genome-wide Survey of the Prevalence and Evolutionary Forces Acting on Human Nonsense SNPs. Am. J. Hum. Genet. 84: 224–234.

